# PACAPergic BAC engages basolateral amygdalar and anterodorsal thalamic networks during looming-threat learning and memory in a labyrinth

**DOI:** 10.64898/2026.06.30.735686

**Authors:** Limei Zhang, Vito S. Hernández, Hai-ying Zhang, Sunny Z. Jiang, Pedro Segura-Chama, Lee E. Eiden

## Abstract

The bed nucleus of the anterior commissure (BAC) is a small pituitary adenylate cyclase-activating polypeptide (PACAP)-rich glutamatergic cell population located at the intersection of the anterior commissure, stria terminalis, stria medullaris, and fornix. Here, we provide functional neuroanatomical characterization of the BAC using PACAP-Cre mice and Cre-dependent viral tracing. BAC axons traverse these major forebrain conduits to engage distributed limbic and diencephalic networks, with dense innervation of the posterior basolateral amygdala (pBLA) and anterior dorsal thalamic nucleus (AD), both implicated in emotional processing and spatial orientation. To determine the functional significance of the BAC, we developed a novel looming-threat memory test (LTMT) paradigm in which mice learned the location of a shelter within a dual-maze labyrinth before exposure to an overhead looming stimulus. Chemogenetic inhibition of BAC PACAP neurons impaired shelter-directed escape, increased freezing behavior, prolonged escape trajectories, and disrupted efficient safety-seeking responses. Selective deletion of PACAP from BAC neurons also blocked key aspects of the looming-induced behavioral phenotype. Fos mapping revealed robust neuronal activation by the looming stimulus, within the AD, pBLA, and dorsal periaqueductal gray (dPAG). BAC silencing blocked Fos induction. PACAP^BAC^ depletion blocked fos induction in AD, but not in pBLA or dPAG. Preliminary *ex-vivo* recordings further indicated that PACAP modulates the intrinsic excitability of AD (and pBLA) neurons. Together, these findings identify the BAC as a previously unrecognized PACAPergic forebrain hub that links emotional and spatial-orientation networks to coordinate looming-threat learning, memory, and adaptive defensive behavior.

## Introduction

Successful defensive behavior requires more than the rapid detection of danger. Animals confronted with an immediate threat must integrate emotional evaluation with spatial memory to generate efficient goal-directed escape, including assessment of locations of safety. This transformation from threat perception to adaptive action recruits distributed neural circuits that include the amygdala, hippocampal formation, anterior thalamus, hypothalamus, and periaqueductal gray (^1^) (^2^). While the individual contributions of these structures to fear, navigation, and defensive responses have been extensively investigated, considerably less is known about the forebrain nodes that coordinate communication among these functional systems.

Pituitary adenylate cyclase-activating polypeptide (PACAP) is a highly conserved neuropeptide that functions as both a neurotransmitter and neuromodulator throughout the mammalian brain (^3–6^). PACAP signaling is critically involved in stress adaptation, arousal, anxiety, emotional learning, and neuroendocrine responses to threat (put Zhang et al., 2021 and Jiang et al., 2023 here). Dysfunction of the PACAP/PAC1 receptor system has been implicated in stress-related psychiatric disorders (^7, 8^). Although PACAP has been studied extensively in the hypothalamus, amygdala, and brainstem, the physiological significance of many discrete forebrain PACAPergic nuclei remains largely unexplored.

The bed nucleus of the anterior commissure (BAC) (^9^), is a small cell population located at the intersection of the anterior commissure, stria terminalis, stria medullaris, and fornix. Historically, the BAC has received remarkably little attention since its original anatomical description nearly a century ago (^9^), and its role in defensive behaviors, including escape, has therefore not been examined in this context. However, recent molecular mapping has revealed that the BAC contains a dense population of PACAP-expressing glutamatergic neurons (^10^), suggesting that it may represent a specialized integrative node embedded within major forebrain fiber systems, and providing a molecular means to assess this small nucleus genetically.

To investigate the BAC by these means, we first characterized the neurochemical identity and long-range projections of BAC PACAP neurons using Cre-dependent viral tracing in PACAP-Cre mice. We then developed a novel *looming-threat memory test* (LTMT) in which mice first learned the location of a shelter within a dual-maze labyrinth before exposure to an overhead looming stimulus, allowing simultaneous assessment of emotional responses and memory-guided navigation. Unlike conventional looming paradigms that evaluate innate escape responses in a familiar open arena, the LTMT requires animals to retrieve previously acquired spatial information to reach safety during an acute visual threat. We then combined chemogenetic inhibition, conditional deletion of PACAP, activity-dependent Fos mapping, and *ex-vivo* electrophysiology to determine whether BAC^PACAP^ neurons play a role. in adaptive defensive behavior.

Together, our findings identify the BAC as a previously unrecognized PACAPergic forebrain hub that coordinates emotional and spatial-orientation networks required for memory-guided escape.

## 2. Materials and Methods

Adult male and female PACAP-Cre and PACAP^flox/flox^ mice (8–14 weeks, C57BL/6J background) were used. All procedures were conducted in accordance with institutional and national guidelines for the care and use of laboratory animals (approved by the NIMH Animal Care and Use Committee, ACUC protocol LCMR-08, and UNAM-FM CIEFM-079-2020. Animal allocation. A summary of animals used is provided in Supplemental Information (SI), extended Material and Methods, Table 1.

### 2.1 Anatomical mapping of BAC–PACAP circuitry

PACAP-Cre mice received unilateral AAV-DIO-Syn-hM4D-tdTomato injections into the bed nucleus of the anterior commissure (BAC) (AP +0.02 mm, ML ±0.5 mm, DV –4.0 mm). Four weeks later behavioral experimentation (see below) was carried out and brain tissue was subsequently collected for histological examination to confirm injection siting and visualize BAC neurons and their projections. Projection patterns were visualized in 70 μm brain sections. For more details see SI extended Material and Methods.

### 2.2. Silencing of BAC–PACAP neurons

was carrying out by treatment, of PACAP-Cre mice injected bilaterally four weeks prior in BAC with AAV-DIO-Syn-hM4D-tdTomato, with CNO (1–3 mg/kg, i.p.) 30 min before behavioral assessment. Wild-type littermates served as CNO controls.

### 2.3. PACAP deletion from BAC neurons

was carried out by bilateral BAC injection of AAV-Cre-IRES-Following a four-week recovery period after viral injections, mice were placed individually in the LTMT apparatus and allowed to freely explore the environment for 5 min (training phase). During this period, animals could discover and repeatedly visit the shelter. After minute 5 of exploration (test phase), a looming stimulus was automatically triggered when the mouse entered a predefined virtual trigger zone ("Zone 0") located near the entrance of Maze A. The stimulus consisted of ten repetitions of an expanding black disk (0.5 s) alternating with a TdTomato in PACAP^flox/flox^ mice.

### 2.4. Looming-threat memory test (LTMT)

Following a four-week recovery from surgery and protein expression from instilled AAV vectors in some groups, mice were exposed to a custom looming-threat memory test (LTMT) labyrinth (approved by NIMH-ACUC protocol LCMR-08) (Fig. 1).

**Figure 1.**
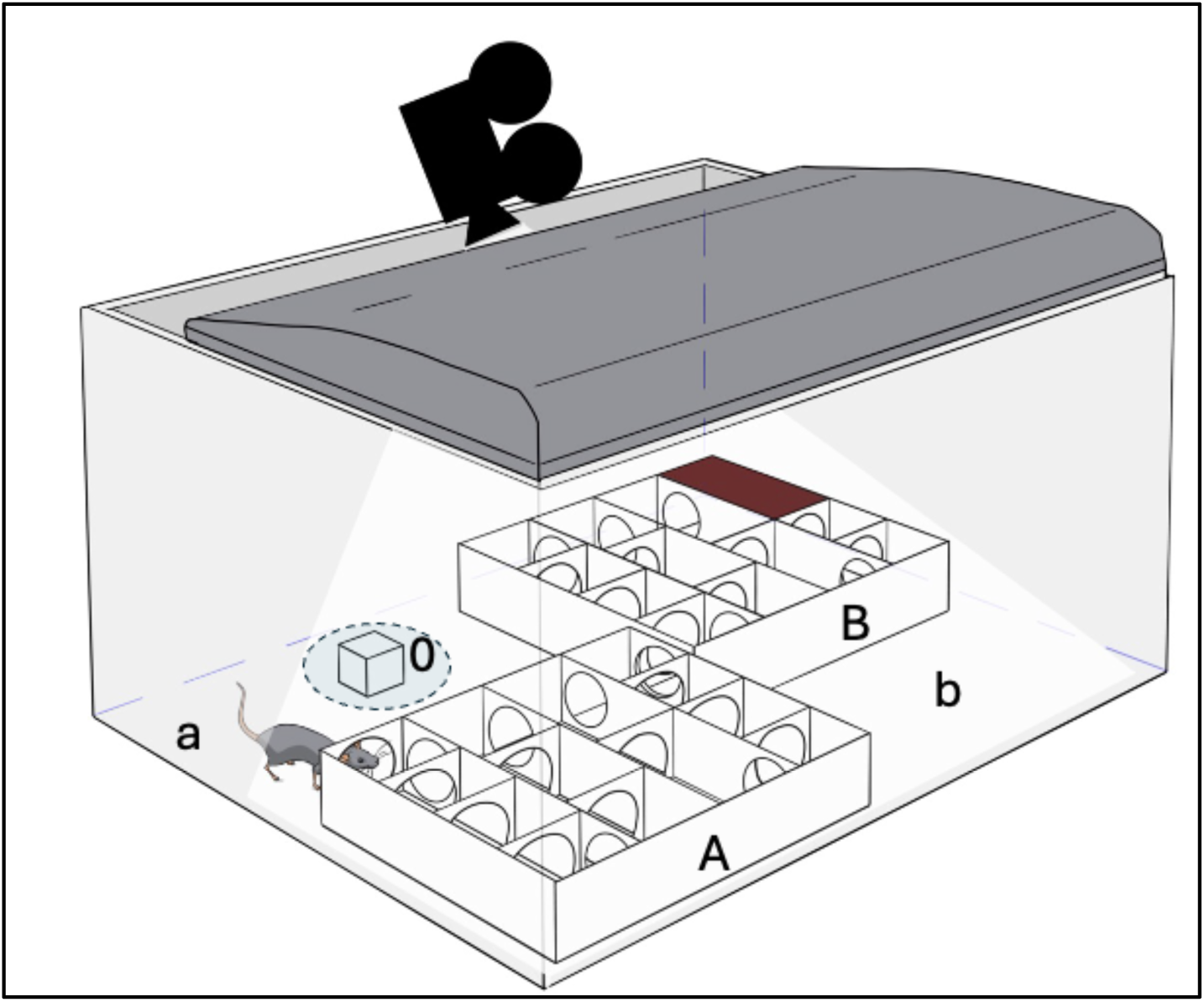
Looming-threat memory test (LTMT). Schematic illustration of the custom behavioral testing apparatus developed for this study. The apparatus consisted of multiple compartments linked via circular tunnels in a maze configuration within a 50 cm x 50 cm x 50 cm standard looming box containing two wooden multi-chamber mazes 24.3 cm x 24.3 cm x 4 cm, with four corridors connected by circular tunnels. The wooden mazes (Wooden Multi-Compartment Hideaway Tunnel Maze, Temu.com) were treated with sealant by the Section on Instrumentation (SOI, NIMH) for suitability for decontamination in accordance with NIMH guidelines. The two mazes (A and B, Fig. 1) are connected in Z form. Outside of A, there is a 24 cm x 24 cm space, "a", in which a plastic cube, 2 x 2 x 2 (cm^3^) with uneven surfaces was placed. A digitally designated 5 cm diameter area within this space was assigned as area "0". Maze B contains a shelter at the far corner occupying two cubic spaces with a connecting tunnel. The shelter has a dark, infrared-transparent plexiglass ceiling (symbolized with dark brown rectangle). A flavored/hydrated gel was placed inside the shelter, which provides episodic and spatial memory of a safe and comfortable location and food reward. Mice first freely explored the dual-maze labyrinth to learn the location of the sheltered compartment for 5 mins. The digital circle "0" (virtual trigger zone) was activated at minute 6 and the test phase, in which a looming visual stimulus was automatically triggered, began when the animal entered zone 0. Escape latency, trajectory, freezing behavior, and time spent in the shelter were recorded (infrared sensitive camera symbolized).

Behavioral assessment was performed using a novel *LTMT* developed for this study (Fig. 1). The apparatus consisted of two interconnected multi-compartment mazes (A and B) forming a labyrinth placed within a standard looming box. Maze B contained a sheltered compartment located at the most distant corner of the apparatus and equipped with a dark infrared-transparent ceiling to permit behavioral tracking. A hydrated flavored gel was placed inside the shelter to encourage exploration and facilitate the formation of spatial memory for a safe location.

Behavior was recorded using automated video tracking software. Primary outcome measures included escape latency, escape trajectory length, velocity, freezing episodes, freezing duration, and time spent inside the shelter. Sixty minutes after behavioral testing, animals were perfused for Fos immunohistochemistry.

Complete apparatus specifications, behavioral acquisition parameters, exclusion criteria, and sanitation procedures are provided in the Supplementary Information, extended Material and Methods.

### 2.5 Ex-vivo whole-cell electrophysiology

Detailed procedures are described elsewhere (^11^). Full methodical description, slicing solutions, recording conditions, and statistical details are provided in SI extended Materials and Methods.

### 2.6 Data

were analyzed using parametric statistics (t-tests or ANOVA; α = 0.05).

## 3. Results

### 3.1 The bed nucleus of the anterior commissure is a distinct PACAP-rich glutamatergic forebrain nucleus

The bed nucleus of the anterior commissure (BAC) is a small forebrain cell group located at the intersection of the anterior commissure, stria terminalis, stria medullaris, and fornix. Although the BAC has been recognized anatomically for approximately one century (^9^), and appears in classical descriptions of the posterior septal region, it has received remarkably little functional attention. Previous studies of the septo-habenular system have focused primarily on the triangular septum and septofimbrial nucleus, and thus the BAC has largely remained an anatomically defined but functionally unexplored nucleus.

Here, we first summarize the previously established chemoanatomical organization of the BAC (^10^). BAC neurons densely express *Adcyap1* (PACAP) and predominantly co-express the glutamatergic marker *Slc17a6* (VGluT2), exhibit weaker *Slc17a7* (VGluT1) expression, express *Calb2* (calretinin), and show little overlap with the GABAergic marker *Slc32a1* (VGAT) (Fig. 2). These observations identify the BAC as a distinct PACAPergic glutamatergic nucleus strategically positioned among major forebrain fiber systems and provide the anatomical rationale for investigating its functional role in coordinating emotional and spatially directed defensive behavior.

**Figure 2.**
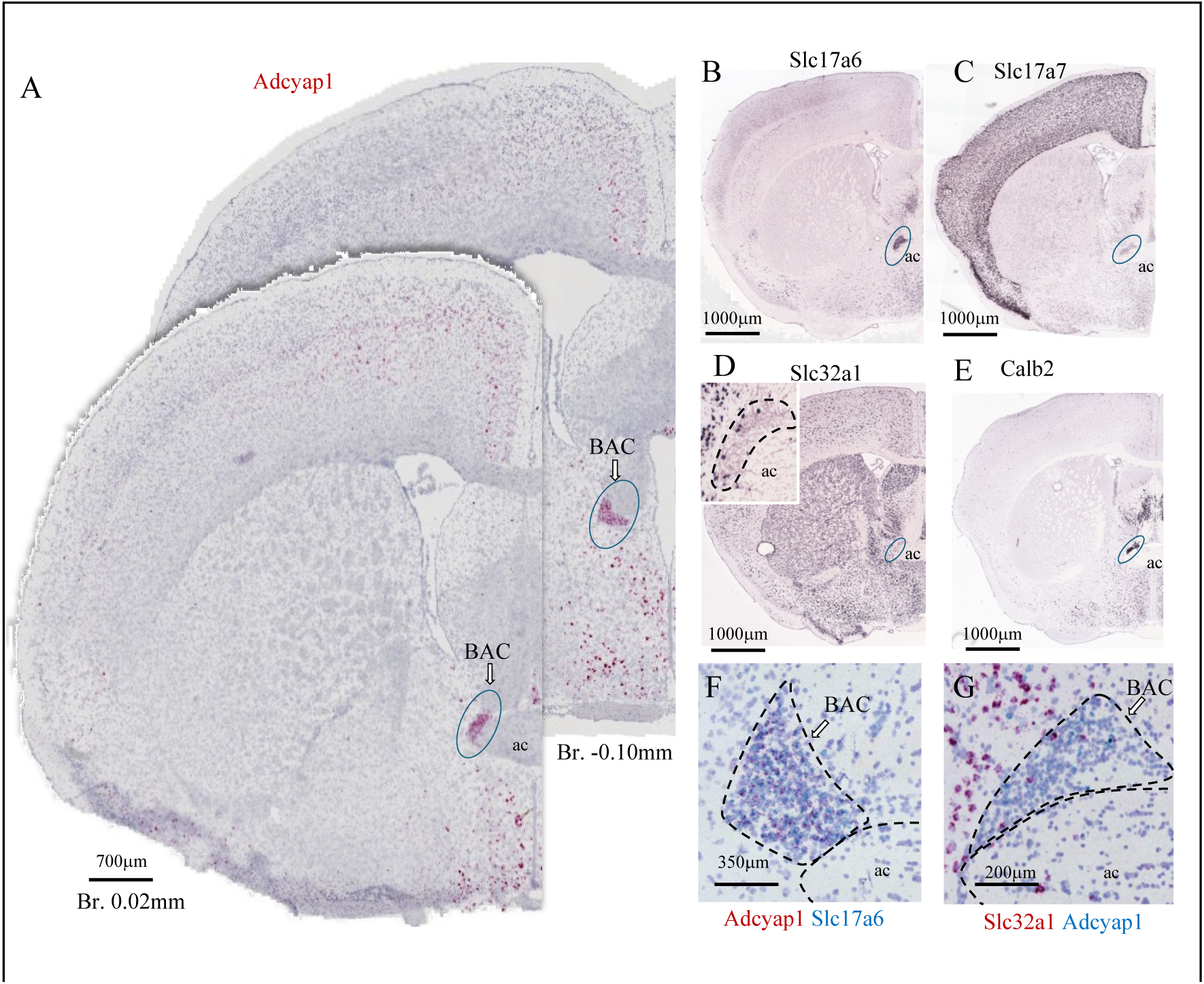
Chemoanatomical organization of the bed nucleus of the anterior commissure (BAC). (A) RNAscope *in situ* hybridization demonstrating dense *Adcyap1* (PACAP) expression in the BAC in two coronal mouse brain sections (Bregma +0.02 mm and −0.10 mm). (B–E) Representative Allen Brain Atlas images showing expression of *Slc17a6* (VGluT2), *Slc17a7* (VGluT1), *Slc32a1* (VGAT), and *Calb2* (calretinin) within the BAC region. (F) Dual *in situ* hybridization demonstrates extensive co-expression of *Adcyap1* and *Slc17a6*, whereas (G) *Adcyap1* and *Slc32a1* show little overlap, indicating that BAC PACAP neurons are predominantly glutamatergic and largely distinct from neighboring GABAergic populations. This figure is reproduced and adapted from Zhang et al., *eLife* (^10^), under the Creative Commons Attribution License (CC BY 4.0), and is included here to provide the anatomical framework for the present functional study.

### 3.2 BAC PACAP neurons engage major forebrain communication pathways and preferentially innervate basolateral amygdalar and anterior dorsal thalamic networks

Since the BAC is positioned at the intersection of the anterior commissure, stria terminalis, stria medullaris, fornix, and medial forebrain bundle, we hypothesized that it distributes information through existing forebrain communication pathways rather than via a single dedicated projection. To examine this possibility, we took advantage of the observation that PACAP expression is a specific marker for BAC neurons, and performed Cre-dependent viral tracing following stereotaxic injections into the BAC of PACAP-Cre mice.

Labeled axons exited the BAC through all five major fiber systems, with particularly prominent labeling within the stria terminalis and stria medullaris (Fig. 3 A,B). These conduits distributed BAC projections throughout multiple limbic and diencephalic structures, including the lateral septum, paraventricular thalamus, lateral habenula, medial amygdala, posterior hypothalamus, and mammillary region (Fig. 3 C–F). For a comprehensive description, see the SI Table 1.

**Figure 3.**
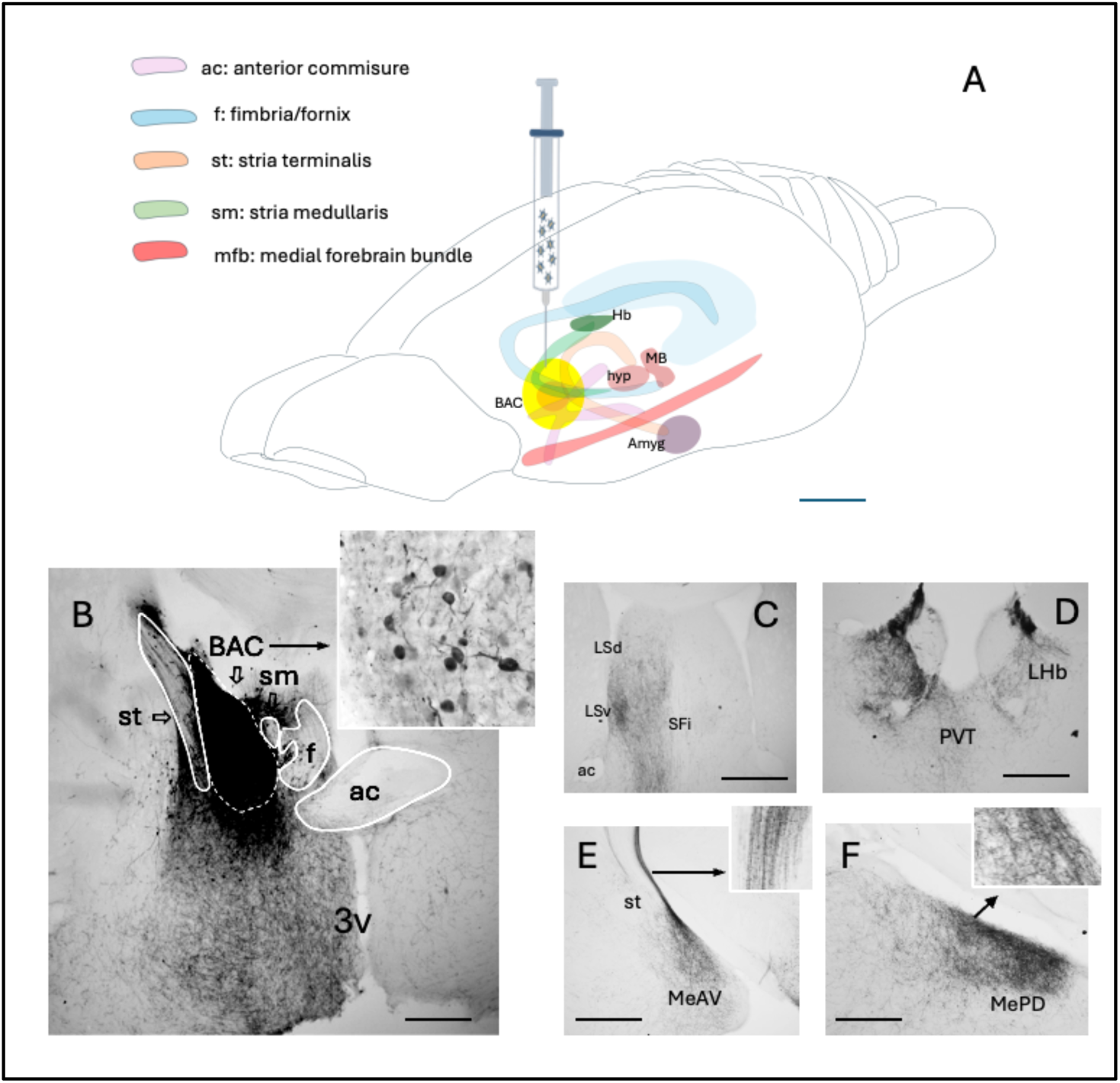
BAC PACAP neurons distribute axons through major forebrain fiber systems. (A) Schematic representation illustrating the anatomical position of the BAC at the intersection of the anterior commissure (ac), fornix (f), stria terminalis (st), stria medullaris (sm), and medial forebrain bundle (mfb), together with the principal projection pathways identified following Cre-dependent viral tracing. (B) Representative injection site within the BAC and labeled fibers entering the major forebrain conduits. Insets show the injection core and labeled axons within the stria terminalis. (C–F) Representative examples of BAC terminal fields in the lateral septum, paraventricular thalamus/lateral habenula, medial amygdala, and posterodorsal medial amygdala. BAC PACAP neurons therefore access multiple limbic and diencephalic structures through well-characterized forebrain communication pathways.

Among all projection fields, two regions consistently exhibited particularly dense terminal innervation: the anterior dorsal thalamic nucleus (AD) and the posterior basolateral amygdala (pBLA) (Fig. 4). Dense terminal arborizations occupied much of the AD, while particularly intense labeling was observed in the posterior division of the basolateral amygdala compared with more anterior regions.

**Figure 4.**
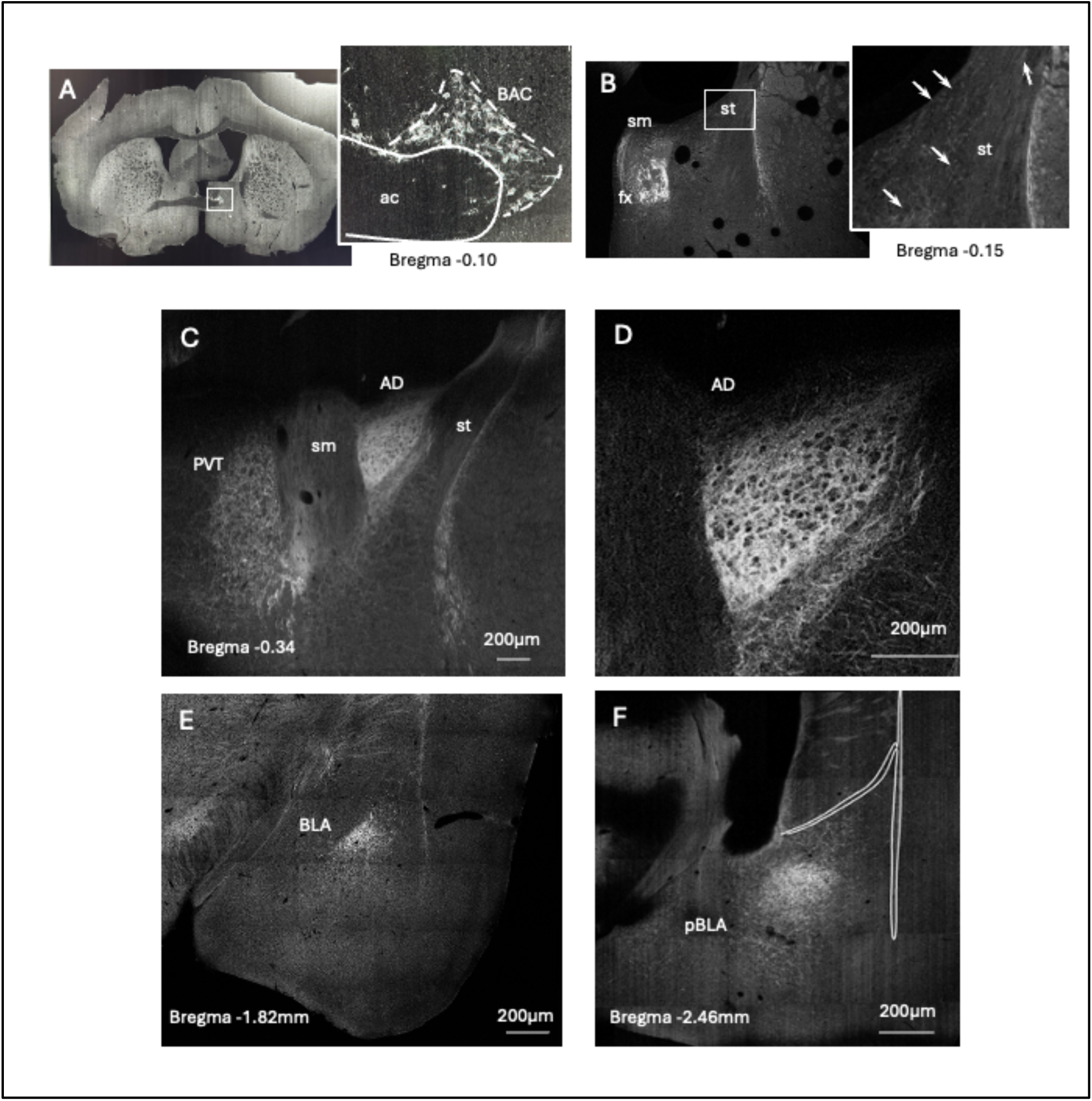
The anterior dorsal thalamic nucleus and posterior basolateral amygdala are major downstream targets of BAC PACAP neurons. (A) BAC injection site. (B) Labeled axons exit the BAC through the stria terminalis (st) and stria medullaris (sm). (C,D) Dense terminal arborization within the anterior dorsal thalamic nucleus (AD). (E,F) BAC fibers innervate the basolateral amygdala, with particularly dense labeling in its posterior division (pBLA). These observations identify AD and pBLA as two principal downstream targets selected for subsequent physiological and functional analyses.

Together, these observations indicate that BAC PACAP neurons are anatomically positioned to coordinate distributed forebrain networks by engaging major white matter communication pathways while preferentially influencing anterior dorsal thalamic and posterior basolateral amygdalar circuits.

### 3.3. PACAP directly modulates the intrinsic excitability of principal BAC target neurons

To determine whether PACAP directly modulates the physiological properties of a major BAC target, whole-cell current-clamp recordings were performed in acute coronal brain slices containing the anterior dorsal thalamic nucleus (AD). Bath application of PACAP (100 nM) revealed two electrophysiologically distinct neuronal responses (Fig. 5).

**Figure 5.**
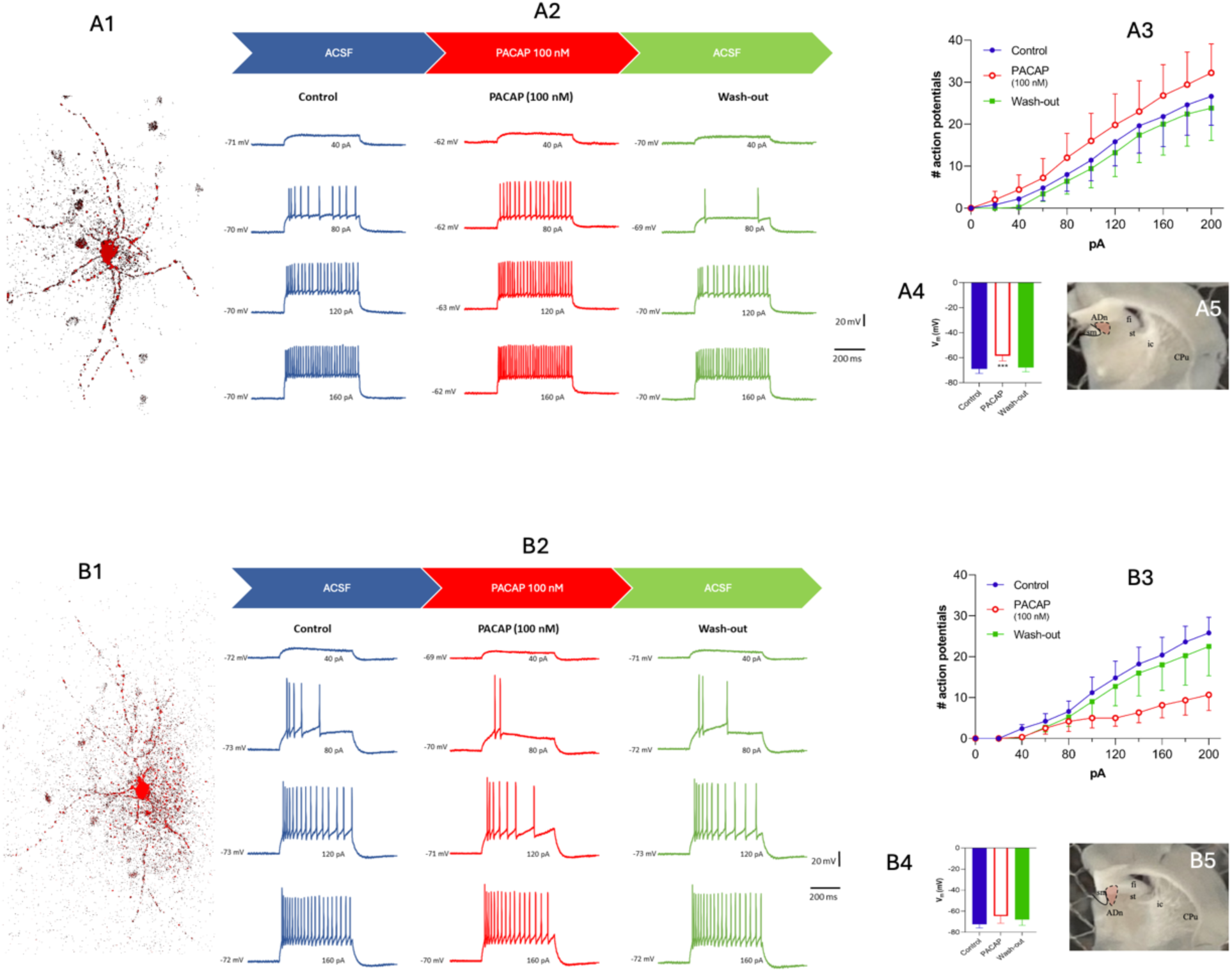
PACAP modulates the intrinsic excitability of anterior dorsal thalamic neurons. Representative whole-cell current-clamp recordings from anterior dorsal thalamic (AD) neurons during baseline artificial cerebrospinal fluid (ACSF), PACAP (100 nM), and washout conditions. (A) Example of a PACAP-responsive neuron showing membrane depolarization and increased firing frequency during PACAP application, together with its reconstructed morphology, current-frequency (I–F) relationship, resting membrane potential measurements, and recording location. (B) Example of a second neuronal population exhibiting reduced excitability during PACAP exposure. These observations demonstrate that PACAP directly modulates the intrinsic electrophysiological properties of AD neurons, supporting the functional responsiveness of a principal downstream target of BAC PACAP neurons.

A first population of AD neurons (PACAP-excited AD neuros, Type I; *n* = 5) exhibited membrane depolarization from −69.0 ± 3.5 mV to −58.6 ± 4.0 mV (*P* = 0.0004, paired *t*-test), accompanied by a significant increase in firing frequency (mixed-effects model; fixed effect of treatment: F(2,20) = 46.0, *p < 0.0001*). Input resistance remained unchanged, indicating that the increase in excitability was not associated with a measurable change in passive membrane properties.

In contrast, a second population of neurons (PACAP-inhibited AD neuros, Type II; *n* = 5) showed a marked reduction in firing frequency during PACAP application (mixed-effects model; fixed effect of treatment: F(2,20) = 13.0, *p = 0.0002*). Resting membrane potential changed only modestly (−72.4 ± 3.4 mV in control versus −64.6 ± 6.8 mV during PACAP application), and input resistance remained unchanged.

In both neuronal populations, the effects of PACAP were reversible after approximately 15 min of washout, indicating an acute modulatory action of the peptide. These observations demonstrate that PACAP directly regulates the intrinsic excitability of AD neurons, supporting the physiological responsiveness of one of the principal downstream targets of BAC PACAP neurons.

### 3.4. BAC PACAP signaling is required for memory-guided defensive behavior

To determine the functional significance of BAC PACAP neurons, animals were tested using the looming-threat memory test (LTMT) following chemogenetic inhibition of BAC neurons or selective deletion of PACAP within the BAC. Under control conditions, mice rapidly navigated toward the previously learned shelter using efficient trajectories and remained inside the shelter after escape (Fig. 6A).

**Figure 6.**
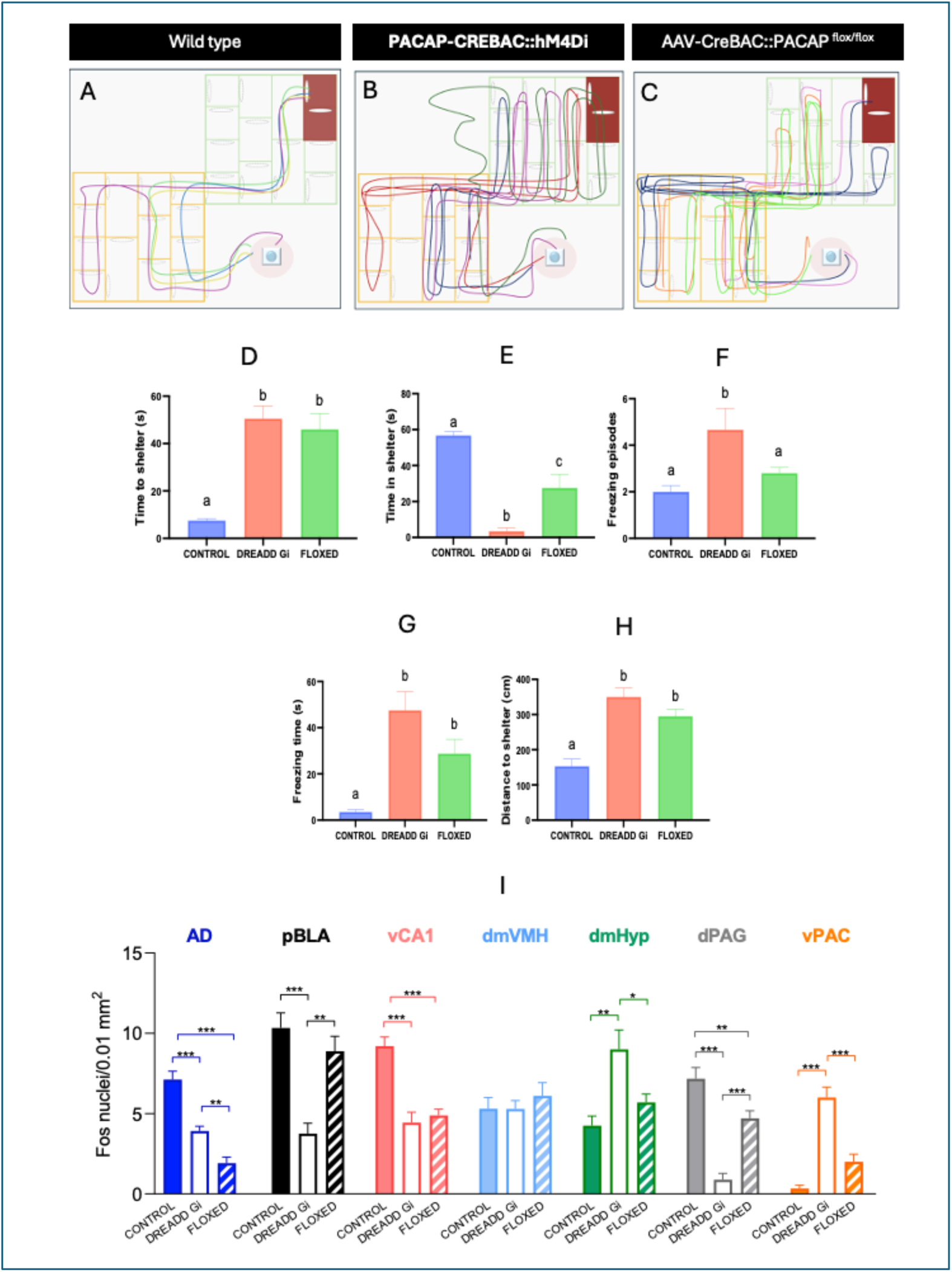
BAC PACAP signaling coordinates memory-guided defensive behavior and downstream circuit recruitment during the looming-threat memory test (LTMT). (A–C) Representative escape trajectories of control mice, mice expressing inhibitory hM4Di DREADDs in BAC PACAP neurons, and mice with selective PACAP deletion in the BAC. Control animals rapidly navigated toward the shelter, whereas BAC inhibition or PACAP deletion produced longer escape trajectories with increased freezing (D–H). Quantification of escape latency, time spent inside the shelter, freezing episodes, freezing duration, and escape path length. Both BAC inhibition and PACAP deletion significantly impaired memory-guided escape behavior. (I) Fos immunohistochemistry performed 60 min after behavioral testing revealed altered recruitment of downstream networks. BAC manipulation significantly reduced neuronal activation within the anterior dorsal thalamic nucleus (AD), posterior basolateral amygdala (pBLA), dorsal periaqueductal gray (dPAG), and ventral PACAPergic hypothalamic region (vPAC), while ventral hippocampal CA1 showed a selective reduction following BAC inhibition. These findings indicate that BAC PACAP neurons coordinate distributed emotional and spatial-defense networks during adaptive responses to looming threat. See SI table 1 for details experimental subjects description.

Both BAC inhibition and selective PACAP deletion markedly impaired defensive performance. Animals required significantly longer to reach the shelter, followed longer and less efficient escape trajectories, exhibited increased freezing episodes and freezing duration, and spent less time inside the shelter following escape (Fig. 6, B–H). These findings indicate that BAC PACAP signaling is required for the efficient transformation of threat detection into memory-guided defensive navigation.

To identify downstream networks associated with these behavioral deficits, Fos expression was quantified in major BAC projection targets and defensive-control regions. BAC inhibition significantly reduced neuronal activation within the anterior dorsal thalamic nucleus (AD), posterior basolateral amygdala (pBLA), dorsal periaqueductal gray (dPAG) and ventral hippocampal CA1. In contrast, Fos levels in vPAC and dmHyp were increased with BAC silencing, while the dorsomedial ventromedial hypothalamus exhibited no changes.

## Discussion

The present study provides the first functional characterization of the bed nucleus of the anterior commissure (BAC), a small but highly distinctive PACAPergic glutamatergic nucleus located at the intersection of several major forebrain fiber systems. Although the BAC has been recognized anatomically since the early neuroanatomical studies of Gurdjian (1927) (^9^), remarkably little has been learned about its function within the brain. By combining viral tracing, chemogenetic manipulation, conditional PACAP deletion, Fos mapping, and *ex-vivo* electrophysiology, we demonstrate that the BAC is positioned to coordinate emotional and spatial-orientation networks required for adaptive defensive behavior.

Our anatomical observations indicate that the BAC occupies a privileged anatomical position embedded within the anterior commissure, stria terminalis, stria medullaris, fornix, and medial forebrain bundle. Instead of projecting selectively to a single downstream structure, BAC PACAP neurons distribute information through these established communication pathways to multiple limbic and diencephalic regions. Among these projections, the particularly dense innervation of the posterior basolateral amygdala and anterior dorsal thalamic nucleus is especially noteworthy. The basolateral amygdala has long been recognized as a critical structure for emotional learning and threat evaluation (^1, 12^), whereas the anterior dorsal thalamic nucleus forms part of the head-direction and spatial orientation system that interacts closely with retrosplenial and hippocampal networks (^13–16^). The preferential innervation of these two regions suggested that the BAC may coordinate emotional and spatial components of defensive behavior.

To test this possibility, we developed the looming-threat memory test (LTMT), a behavioral paradigm that differs from conventional looming assays by requiring animals to retrieve previously acquired spatial information before initiating escape. Most looming paradigms examine innate defensive responses in open arenas (^17^). In contrast, the LTMT requires successful integration of threat detection with memory-guided navigation toward a previously learned shelter. BAC inhibition or selective deletion of PACAP disrupted this process, producing longer escape trajectories, delayed shelter seeking, and increased freezing. These observations indicate that BAC PACAP neurons contribute not only to emotional responses to threat but also to the efficient transformation of remembered spatial information into adaptive defensive action.

The Fos mapping further supports this interpretation. Reduced activation of the posterior basolateral amygdala following BAC manipulation is consistent with impaired recruitment of emotional processing circuits, whereas decreased Fos within the anterior dorsal thalamic nucleus suggests reduced engagement of thalamic spatial-orientation networks. Altered activity in the dorsal periaqueductal gray and ventral PACAPergic hypothalamic regions further indicates that BAC influences downstream motor and autonomic components of defensive behavior. Although these observations do not establish direct functional connectivity, they define a distributed network whose coordinated recruitment depends upon intact BAC PACAP signaling. The observation that PACAP directly modulates the intrinsic excitability of anterior dorsal thalamic neurons further supports the physiological relevance of this pathway. (References)

Our findings also extend recent work identifying the anterior dorsal thalamic nucleus (AD) as a dynamic integrative component of the head-direction network rather than a passive relay of spatial information. Recent physiological studies have shown that AD neurons continuously integrate multimodal sensory signals, behavioral state, and movement to update orientation during navigation (^15, 16^), while pathological disruption of AD activity impairs directional coding and produces spatial disorientation during memory acquisition (^18^). Our observations that BAC inhibition markedly reduced Fos activation in the AD, disrupted efficient shelter-directed trajectories, and prolonged escape despite previous learning of the shelter location are consistent with this emerging framework. Rather than suggesting a primary deficit in spatial memory formation, these findings support the interpretation that BAC PACAP neurons recruit anterior thalamic circuits required for the rapid retrieval and updating of directional information under threatening conditions. We suggest that the BAC acts upstream of the AD to couple emotionally salient sensory information with thalamic spatial-orientation networks, enabling efficient memory-guided defensive escape.

Some limitations should be acknowledged. The present study was designed to identify the BAC as a functional node within larger defensive networks rather than to establish synaptic mechanisms in each downstream target. The electrophysiological experiments presented here provide evidence that PACAP directly influences the excitability of principal BAC target neurons but do not determine the endogenous mode of peptide release during behavior. Likewise, Fos mapping identifies candidate downstream regions but cannot distinguish direct from polysynaptic circuit effects. Future studies combining optogenetics, projection-specific manipulations, and simultaneous recordings will be required to define the dynamic interactions among the basolateral amygdala, anterior dorsal thalamic nucleus, hippocampus, and associated cortical regions during memory-guided defensive behavior.

In summary, our findings identify the BAC as a previously unrecognized PACAPergic forebrain hub that links emotional evaluation with spatial orientation during adaptive defensive behavior. Rather than functioning as an isolated projection system, the BAC appears to engage distributed communication pathways that integrate limbic and thalamic networks, enabling efficient memory-guided escape in response to acute visual threat.

## Acknowledgments

This work was supported by the Universidad Nacional Autónoma de México, UNAM-PAPIIT (IG200121 to L.Z. and IT200125 to V.S.H.), the Mexican Secretaría de Ciencia, Humanidad, Tecnología e Innovación (SECIHTI): CIORGANISMOS-2025-92 to L.Z.: CF-2023-G-243) and the Intramural Research Program of the National institute of Mental Health, NIMH, NIH (ZIAMH002386 to L.E.E.). The authors acknowledge the PASPA-DGAPA sabbatical fellowships (to L.Z. and V.S.H.), a SECIHTI sabbatical fellowship (to L.Z.), and a Fulbright-García Robles Fellowship (to V.S.H.). We thank the NIMH Rodent Behavioral Core (Director Yogita Chudasama, and staff members Alice Graham, Sean Bradley and Kevin Cravedi), NIMH Section on Instrumentation (Director: George Dodd, staff member Eric Saglio), NIMH Systems Neuroscience Imaging Resource (Director: Ted Usdin, staff members Jonathan Kuo and Vitaly Boyko) for guidance in instrument fabrication and experimental work and Oscar Hernández, Maria José Gómora for technical assistance.

## Supplemental Information

### Supplemental Extended Materials and Methods

#### Animals, experimental design, and allocation

Adult male and female C57BL/6J PACAP-Cre and PACAP^^flox/flox^ mice, 8–14 weeks of age, were used in this study. Animals were housed under standard vivarium conditions with ad libitum access to food and water unless otherwise specified. All procedures were approved by the NIMH Animal Care and Use Committee, protocol LCMR-08, and the UNAM-FM committee, CIEFM-079-2020.

Animals were assigned to experimental groups for anatomical tracing of BAC PACAP neurons, chemogenetic inhibition of BAC PACAP neurons, selective deletion of PACAP from BAC neurons, looming-threat memory testing and Fos mapping, and ex-vivo whole-cell electrophysiology.

Injection sites and viral expression were verified histologically. Animals were excluded if viral expression was absent, if the injection site was outside the BAC, if expression spread extensively into adjacent nuclei. Final animal numbers are summarized in Table S1.

**Table S1.** Animal allocation and experimental cohorts.

| Experiment | Genotype | Viral manipulation | Treatment | Sex | Final N |
| --- | --- | --- | --- | --- | --- |
| BAC anatomical tracing | PACAP-Cre | Unilateral AAV-DIO-Syn-hM4D-tdTomato into BAC | None | 2 M / 1 F | 3 |
| Control LTMT + Fos | Wild-type littermates | Control surgical sham stereotaxic injection | CNO, 3 mg/kg, i.p. | 5 M / 5 F | 10 |
| BAC chemogenetic inhibition | PACAP-Cre | Bilateral AAV-DIO-Syn-hM4D-tdTomato into BAC | CNO, 3 mg/kg, i.p. | 5 M / 5 F | 10 |
| BAC PACAP deletion | PACAP<sup>^flox/flox</sup> | Bilateral AAV-Cre-IRES-dsTomato into BAC | None | 5 M / 5 F | 10 |
| AD electrophysiology | PACAP-Cre or wild-type mice | None | PACAP, 100 nM ex vivo | Male | 10 neurons from 5 mice |

#### Stereotaxic surgery and viral injections

Mice were anesthetized with a 100mg/kg ketamine and 8 mg/kg xilazine i.p. dose and secured in a stereotaxic frame. Body temperature was maintained with a heating pad throughout surgery. After scalp incision and skull leveling, small craniotomies were made above the BAC. For anatomical mapping, PACAP-Cre mice received unilateral injections of AAV-DIO-Syn-hM4D-tdTomato into the BAC using the following coordinates relative to bregma: AP +0.02 mm, ML ±0.5 mm, DV −4.0 mm. For chemogenetic inhibition, PACAP-Cre mice received bilateral BAC injections of AAV-DIO-Syn-hM4D-tdTomato. For selective PACAP deletion, PACAP^^flox/flox^ mice received bilateral BAC injections of AAV-Cre-IRES-tdTomato.

Viral vectors were delivered at a volume of 250nl per side and infusion rate of [100 nL/min] using a Hamilton syringe. Injectors were left in place for 10 after infusion to minimize backflow. Mice recovered for four weeks to allow viral expression before behavioral testing or tissue collection. For chemogenetic experiments, mice expressing hM4Di in BAC PACAP neurons received clozapine-N-oxide, CNO, 3 mg/kg, intraperitoneally, 30 min before behavioral testing. Wild-type littermates receiving CNO served as controls.

#### Histology and anatomical mapping of BAC PACAP circuitry

Four weeks after viral injection, animals were deeply anesthetized and perfused transcardially with 0.9% saline followed by 4% paraformaldehyde. Brains were removed, and sectioned coronally at 70 µm using a Leica VT1200 vibratome.

Sections were mounted or processed as free-floating sections and imaged using a confocal microscope. BAC injection sites were confirmed by tdTomato expression within the anatomical boundaries of the bed nucleus of the anterior commissure. Projection patterns were evaluated throughout the forebrain, diencephalon, hypothalamus, amygdala, and midbrain. Labeled fibers were assigned to major forebrain conduits, including the anterior commissure, fornix/fimbria, stria terminalis, stria medullaris, and medial forebrain bundle. Projection density was scored semi-quantitatively as absent, sparse, moderate, dense, or very dense. Representative projection targets are summarized in Table S2.

#### Looming-threat memory test apparatus

Behavioral testing was performed in a custom looming-threat memory test, LTMT, labyrinth placed inside a standard looming box. The outer box measured 50 × 50 × 50 cm. Inside the box, two wooden multi-compartment mazes, designated Maze A and Maze B, were arranged in a Z-shaped configuration. Each maze measured 24.3 × 24.3 × 4 cm and contained four corridors connected by circular tunnels. The wooden mazes were treated with sealant by the NIMH Section on Instrumentation to permit cleaning and decontamination according to institutional guidelines.

Outside Maze A, a 24 × 24 cm open space was designated as area “a.” A small plastic cube measuring 2 × 2 × 2 cm with uneven surfaces was placed in this region. A digitally defined circular trigger zone, Zone 0, 5 cm in diameter, was positioned within this open area.

Maze B contained the shelter compartment at the far corner of the labyrinth. The shelter occupied two connected cubic spaces and was covered by a dark, infrared-transparent plexiglass ceiling, allowing behavioral tracking by an infrared-sensitive camera while maintaining a protected dark compartment. A flavored hydrated gel was placed inside the shelter to encourage exploration and support the acquisition of spatial and episodic information about the safe location.

#### LTMT behavioral procedure

Following four weeks of recovery after viral surgery, mice were placed individually into the LTMT apparatus. During the training phase, mice were allowed to freely explore the full labyrinth for 5 min. During this period, animals could discover, enter, and repeatedly visit the sheltered compartment.

At minute 6, the test phase began. The virtual trigger zone, Zone 0, was activated, and the looming stimulus was automatically initiated when the animal entered this zone. The looming stimulus consisted of ten repetitions of an expanding black disk presented for 0.5 s, alternating with a bright background presented for 0.5 s. This stimulus was designed to mimic the approach of an aerial predator.

Behavior was recorded using automated video-tracking software. Primary outcome variables included escape latency, time spent in the shelter, number of freezing episodes, total freezing duration, escape trajectory length, velocity, and shelter-directed navigation efficiency. Representative escape trajectories were generated from tracking coordinates.

Sixty minutes after behavioral testing, animals assigned to Fos mapping were deeply anesthetized and perfused for immunohistochemical analysis.

#### LTMT acquisition parameters and behavioral definitions

Behavioral videos were acquired under infrared illumination. Escape latency was defined as the time from looming stimulus onset to first entry into the shelter. Shelter time was defined as the cumulative time spent within the shelter compartment after looming stimulus onset. Escape path length was calculated as the cumulative distance traveled from stimulus onset to first shelter entry. Freezing was defined as immobility lasting at least 1 s with no movement except for respiratory motion. Freezing episodes and total freezing time were quantified during the post-stimulus test interval.

Behavioral scoring was performed manually by two independent researchers blinded to group identity during behavioral quantification whenever possible.

#### LTMT exclusion criteria

Animals were excluded from behavioral analysis if one or more of the following occurred: misplaced viral injection outside the BAC, insufficient viral expression, extensive viral spread into adjacent regions, failure of the tracking system during the looming stimulus or escape period, poor video quality preventing reliable tracking, failure of the looming stimulus to trigger, illness or abnormal locomotor impairment unrelated to experimental manipulation, or procedural complications during surgery.

#### Apparatus sanitation

Between animals, the LTMT apparatus was cleaned with 70% ethanol and allowed to dry completely before the next trial. The shelter, tunnels, open areas, plastic cube, and maze surfaces were wiped thoroughly to remove urine, feces, food residue, and odor cues.

#### Fos immunohistochemistry and quantification

Mice assigned to activity mapping were perfused 60 min after LTMT testing. Brains were collected, and sectioned coronally at 70 µm section thickness. Sections containing the anterior dorsal thalamic nucleus, posterior basolateral amygdala, ventral hippocampal CA1, dorsomedial ventromedial hypothalamus, dorsomedial hypothalamus, dorsal periaqueductal gray, and ventral PACAPergic hypothalamic region were processed for Fos immunohistochemistry.

Sections were incubated with anti-Fos primary antibody (rabbit polyclonal antibody to c-FOS, Cat# RPCA-c-FOS) in TBST for 24h at 4°C, followed by a secondary biotinylated donkey anti rabbit antibody . Sections were mounted and imaged under identical acquisition settings within each region.

Fos-positive nuclei were counted in anatomically matched regions of interest. Data were expressed as Fos-positive nuclei per 0.01 mm². For Figure 6I, quantification was performed using 5 mice per group. Counts from multiple fields in a determined region within each animal were averaged to generate one biological replicate per region per animal. Statistical analyses were performed on animal-level means, not on individual fields. Regions were identified using a mouse brain atlas and matched across groups by rostrocaudal level.

#### Ex-vivo whole-cell electrophysiology

Whole-cell patch-clamp recordings were performed in acute coronal brain slices (300 µm) obtained from male mice (eight-week-old) with vibratome Leica 1200 (Leica Biosystems). Mice were deeply anesthetized (5% isoflurane) and quickly decapitated. The brain was dissected and placed in ice-cold oxygenated (95% O_2_/5% CO_2_) high-sucrose artificial cerebrospinal fluid (ACSF; in mM: 80 NaCl, 3.5 KCl, 4 MgSO_4_, 1.25 NaH_2_PO_4_, 25 NaHCO_3_, 0.5 CaCl_2_, 90 sucrose, 10 glucose pH 7.4 and ∼ 290 mOsm). Slices recovered for at least 1h in oxygenated high-sucrose ACSF solution in a recovery chamber at room temperature until recording. Brain slices were bathed at 2 mL/min with ACSF containing in mM: 126 NaCl, 2,5 KCl, 1.2 MgSO_4_, 1.25 NaH_2_PO_4_, 25 NaHCO_3_, 2 CaCl_2_, 10 glucose pH 7.4 and oxygenated with 95% O_2_/5% CO_2_, ∼ 290-300 mOsm. Recordings were obtained from neurons in the anterior dorsal thalamic nucleus using differential interference contrast infrared (DIC-IR) microscopy (Nikon Eclipse N1) equipped with a X 40 water- immersion objective. Patch pipettes (7-10 MΩ) were pulled from borosilicate glass using a puller P-97 (Sutter Instruments) and filled with an intracellular solution containing (in mM): 140 K-gluconate, 10 NaCl, 0.1 CaCl₂, 5 EGTA, 10 HEPES, .005 biocytin, 2 ATP-Mg, and 1 GTP-Na, the pH was adjusted to 7.2 with KOH, and osmolarity was adjusted ∼290 mOsm. Electrophysiological signals were acquired using an Axoclamp-2B amplifier (Axon Instruments), a Digidata 1440 digitizer, and the pClamp 10.4 software (Molecular Devices). Current-clamp protocols were used to assess intrinsic excitability. Current steps of 500 ms duration, from -160 to 200 pA, Δ 20 pA, were injected from resting membrane potential to generate frequency-current (F-I) curves and obtained the input resistance.

#### Statistical analysis

Comparisons between two groups were performed using paired t-tests, as appropriate. Comparisons among three or more groups were performed using one-way or two-way ANOVA followed by post-hoc multiple-comparison tests. The alpha level was set at 0.05.

Behavioral and Fos data were analyzed using animal-level means as biological replicates. For Fos quantification, multiple sections or fields from the same animal were averaged before statistical testing. Electrophysiological data were analyzed using cell-level measurements with the number of cells and animals reported explicitly. Data are presented as mean ± SEM unless otherwise stated.

## Supplemental Results

**SI Table 2.** Distribution of axonal projections in PACAP-Cre mice transfected with AAV-hM4Di virus injected into the BAC.

| Structure | Abbr. |  |
| --- | --- | --- |
| **Conducting Systems** |  |  |
| anterior commissure | ac | + |
| medial forebrain bundle | mfb | +++ |
| septo-fimbria | sf | ++ |
| stria medullaris | sm | ++++ |
| stria terminalis | st | ++++ |
| **Telencephalon** |  |  |
| **Cerebral cortex** |  |  |
| Clastrum |  | + |
| Endopiriform Nu | EP | + |
| Insular cortex |  | + |
| Piriform cortex | Pir | + |
| **Hippocampal formation** |  |  |
| CA1 (ventral) | CA1v |  |
| CA3 (ventral) | CA3v |  |
| Dentate gyrus (temporal) | DG |  |
| **Extended amygdala** |  |  |
| Anterior amygdala | AA | ++ |
| Bed nucleus of anterior commissure | BAC | ++++ |
| Bed nucleus of stria terminalis anterior | STMA | ++++ |
| Bed nucleus of stria terminalis posterior | STMP | ++++ |
| Basolateral amygdala | BLA | + |
| Basomedial amygdala | BMA | ++ |
| Bed nucleus of stria terminalis intra-amygdala division | STIA | + |
| Cortico-amygdala | CoA | ++ |
| Medial amygdaloid nucleus, postero-dorsal | MePD | +++ |
| Medial amygdaloid nucleus, postero-ventral | MePV | +++ |
| NLOT |  |  |
| Cerebral nuclei |  |  |
| Accumbens nucleus | AcbC | + |
| Nu. Basalis of Meynert | B | ++ |
| Diagonal band nucleus | NDB | ++ |
| Ventral pallidum | VP | ++ |
| Lat. septal n. dorsal | LSD | ++ |
| Lat. septal n. int | LSI | ++ |
| Lat. septal n. ventral | LSV | ++++ |
| Medial septal nucleus | MS | ++ |
| Septo-fimbrial n. | SFi | +++ |
| Septo-hippocampal n. | SHi | ++ |
| Substantia innominata | SI | ++ |
| Triangular septum | TS | +++ |
| Diencephalon |  |  |
| Hypothalamus |  |  |
| Anteroventral periventricular n. | AVPe | ++ |
| Rostral periventricular area | RP3V | ++ |
| Anterior hypothalamic area anterior | AHA | +++ |
| Anterior hypothalamic area posterior | AHp | +++ |
| Arcuate nucleus | Arc | ++ |
| Dorsal hypothalamic | DH | ++ |
| Dorsomedial hypothalamic area | DM | ++ |
| Lat. hypothalamic area | LH | +++ |
| Lat. preoptic area | LPO | +++ |
| Mammillary nucleus | MM | +++ |
| Mammillo tuberal nu. | MTu | +++ |
| Medial preoptic area | MPA | ++++ |
| Median preoptic n. | MnPO | ++ |
| Parastrial nucleus | PS | + |
| Paraventricular hypothalamic n. anterior | PVA | +++ |
| Paraventricular hypothalamic n. magnocellular div. | PVNIm | ++ |
| Paraventricular hypothalamic n. parvocellular div. | PVNmp | ++ |
| Periventricular nu. | Pe | + |
| Premammillary n. ventral | PMV | ++ |
| Posterior hypothalamic area | PH | +++ |
| Retrochiasmatic area | RCh | ++ |
| Septohypothalamic area | SHy | ++++ |
| Suprachiasmatic nu. | SCN | ++ |
| Supraoptic nucleus | SON | + |
| Tuber cinereum area | TC | ++ |
| Ventral tuberomammillar n. | VTM | ++ |
| Ventromedial hypothalamic nucleus | VMH | + |
| Ventrolateral preoptic area | VLPO | ++ |
| Thalamus |  |  |
| Anterodorsal thalamic nucleus | AD | ++ |
| Antero medial thalamic nucleus | AM | + |
| Central medial thalamic nucleus | CM | + |
| Lateral geniculate nu. | LG | ++ |
| Mediodorsal thalamic nucleus | IMD | + |
| Paraventricular thalamic nucleus | PV | +++ |
| Reuniens thalamic nucleus | Re | ++ |
| Rhomboid thalamic nucleus | Rh | ++ |
| Zona incerta | ZI | +++ |
| **Epithalamus** |  |  |
| Lateral habenula | LHb | +++ |
| **Mesencephalon** |  |  |
| Dorsal raphe nucleus | DR | + |
| Interpeduncular n. | IP | + |
| Median raphe n. | MnR | + |
| Periaqueductal gray, lateral | LPAG | ++ |
| Periaqueductal gray, dorsomedial | DMPAG | +++ |
| Substantia nigra | SN |  |
| Superior colliculus,<br>intermediate and deep gray layer | DpG | + |
| Supramammillary n. | SuM | ++ |
| Ventral tegmental area | VTA | ++ |

### Supplemental statistics analysis

Fos after 60 minutes after looming test Ordinary one-way ANOVA

All groups were tested for normality using the D’Agostino-Pearson omnibus and Shapiro-Wilk (W) tests. Alpha level was set at 0.05. Tukey’s multiple comparisons test was used as post hoc test to evaluate differences between groups.

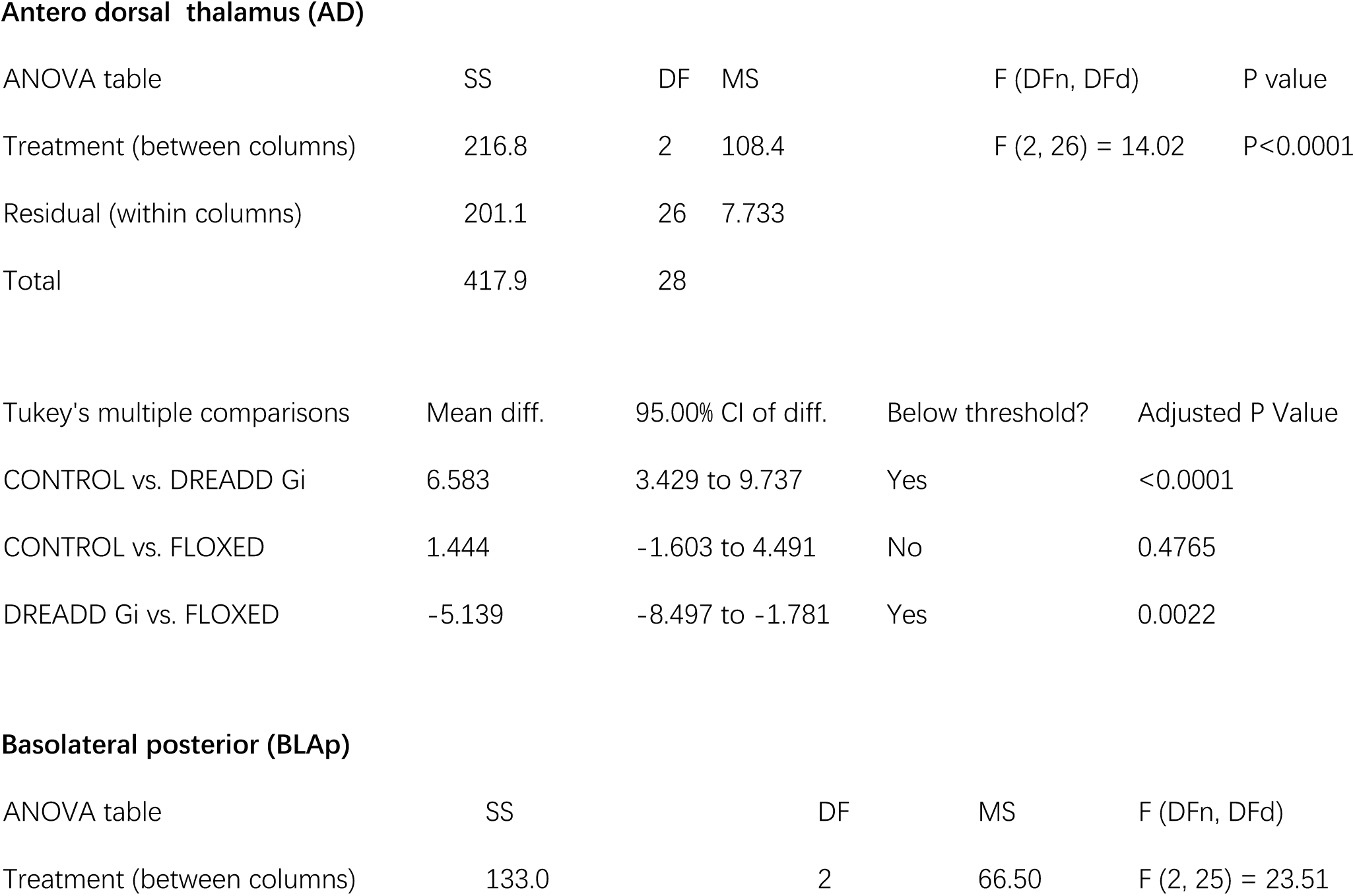
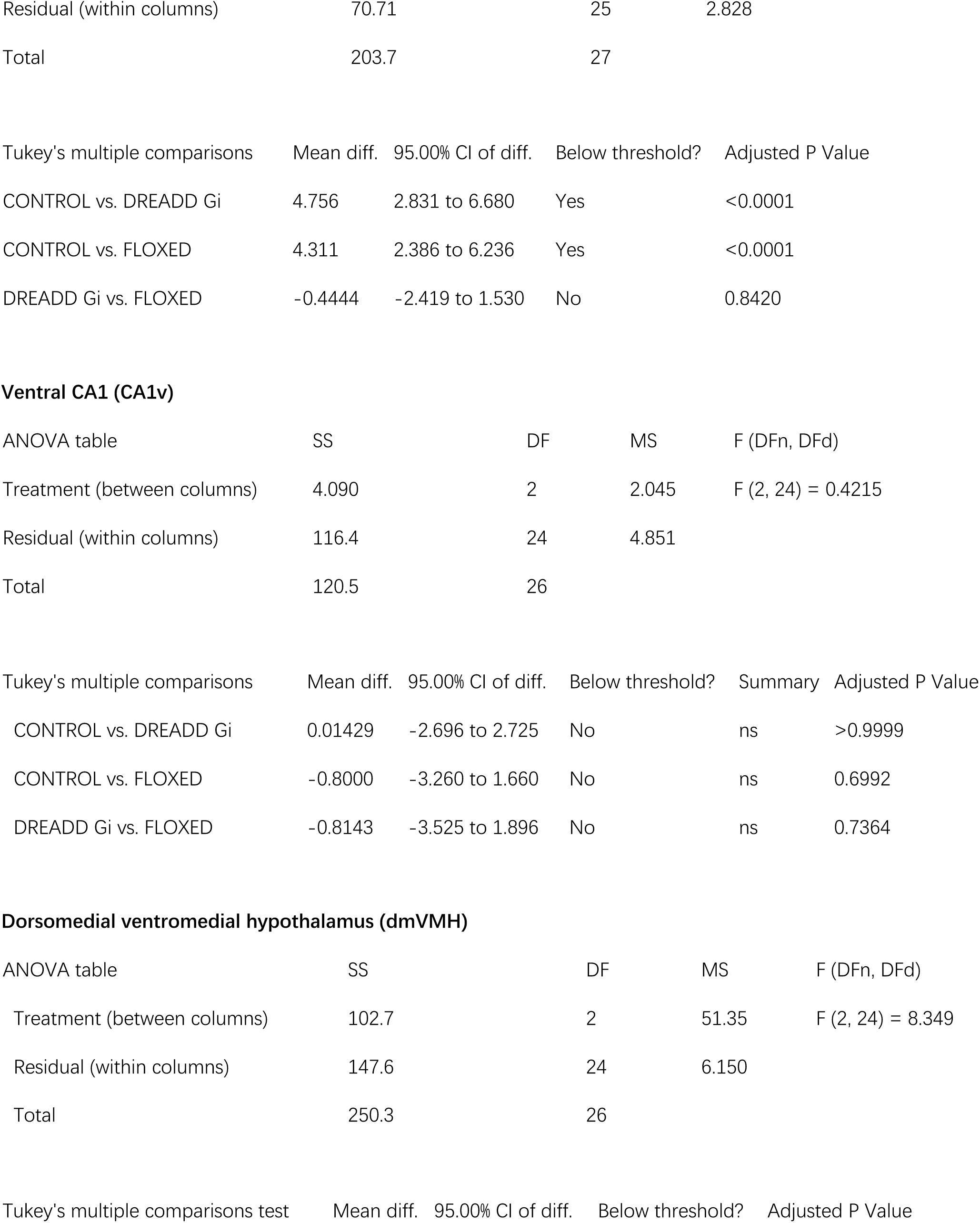
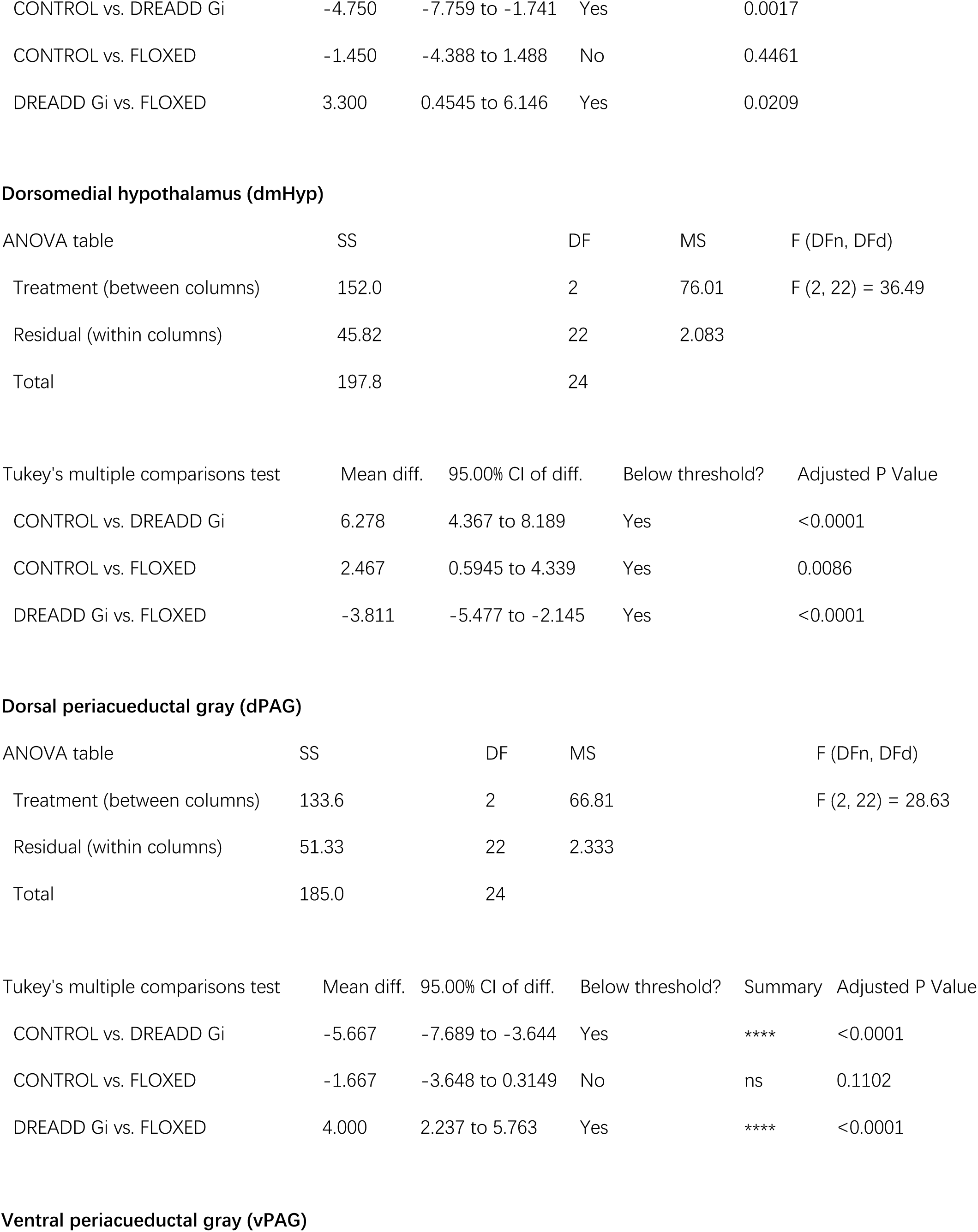
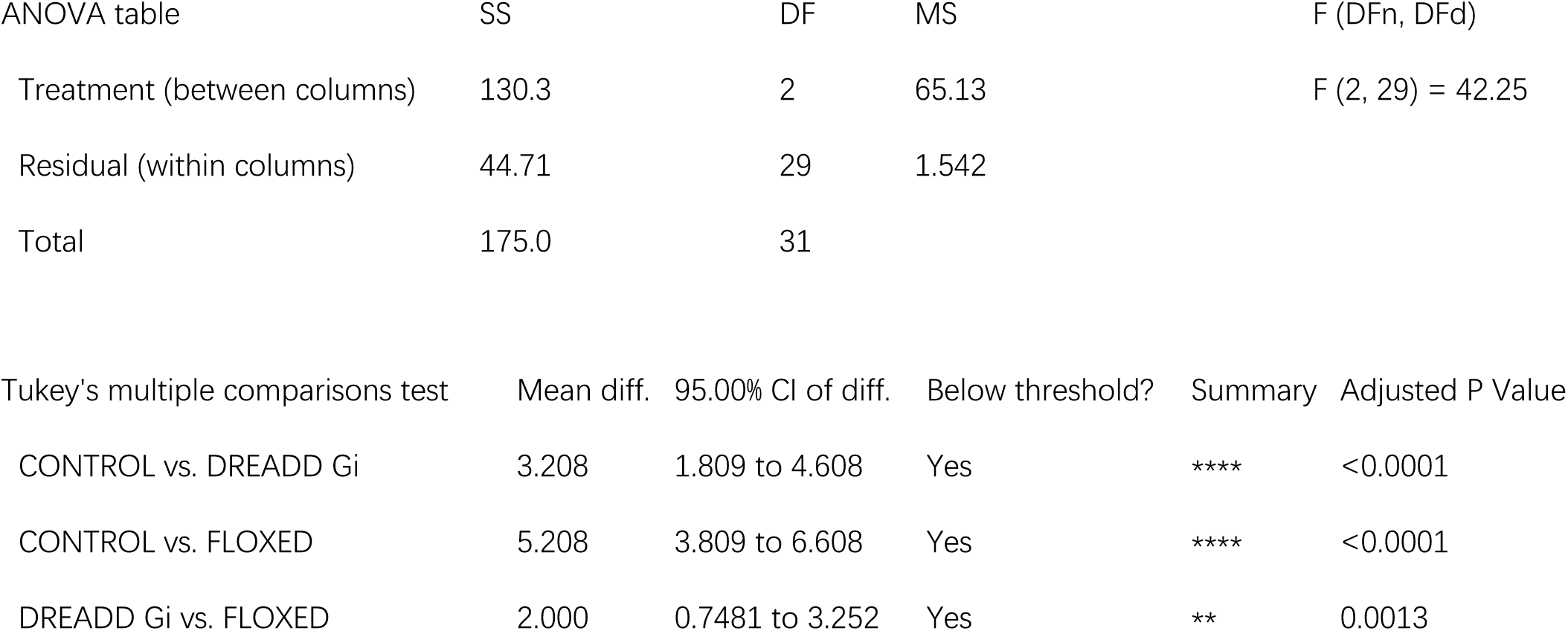

**Antero dorsal thalamus (AD)**
| ANOVA table | SS | DF | MS | F (DFn, DFd) | P value |
| --- | --- | --- | --- | --- | --- |
| Treatment (between columns) | 216.8 | 2 | 108.4 | F (2, 26) = 14.02 | P<0.0001 |
| Residual (within columns) | 201.1 | 26 | 7.733 |  |  |
| Total | 417.9 | 28 |  |  |  |

| Tukey's multiple comparisons | Mean diff. | 95.00% CI of diff. | Below threshold? | Adjusted P Value |
| --- | --- | --- | --- | --- |
| CONTROL vs. DREADD Gi | 6.583 | 3.429 to 9.737 | Yes | <0.0001 |
| CONTROL vs. FLOXED | 1.444 | -1.603 to 4.491 | No | 0.4765 |
| DREADD Gi vs. FLOXED | -5.139 | -8.497 to -1.781 | Yes | 0.0022 |

**Basolateral posterior (BLAp)**
| ANOVA table | SS | DF | MS | F (DFn, DFd) |
| --- | --- | --- | --- | --- |
| Treatment (between columns) | 133.0 | 2 | 66.50 | F (2, 25) = 23.51 |

|  |  |  |  |
| --- | --- | --- | --- |
| Residual (within columns) | 70.71 | 25 | 2.828 |
| Total | 203.7 | 27 |  |

| Tukey's multiple comparisons | Mean diff. | 95.00% CI of diff. | Below threshold? | Adjusted P Value |
| --- | --- | --- | --- | --- |
| CONTROL vs. DREADD Gi | 4.756 | 2.831 to 6.680 | Yes | <0.0001 |
| CONTROL vs. FLOXED | 4.311 | 2.386 to 6.236 | Yes | <0.0001 |
| DREADD Gi vs. FLOXED | -0.4444 | -2.419 to 1.530 | No | 0.8420 |

Ventral CA1 (CA1v)
| ANOVA table | SS | DF | MS | F (DFn, DFd) |
| --- | --- | --- | --- | --- |
| Treatment (between columns) | 4.090 | 2 | 2.045 | F (2, 24) = 0.4215 |
| Residual (within columns) | 116.4 | 24 | 4.851 |  |
| Total | 120.5 | 26 |  |  |

| Tukey's multiple comparisons | Mean diff. | 95.00% CI of diff. | Below threshold? | Summary | Adjusted P Value |
| --- | --- | --- | --- | --- | --- |
| CONTROL vs. DREADD Gi | 0.01429 | -2.696 to 2.725 | No | ns | >0.9999 |
| CONTROL vs. FLOXED | -0.8000 | -3.260 to 1.660 | No | ns | 0.6992 |
| DREADD Gi vs. FLOXED | -0.8143 | -3.525 to 1.896 | No | ns | 0.7364 |

Dorsomedial ventromedial hypothalamus (dmVMH)
| ANOVA table | SS | DF | MS | F (DFn, DFd) |
| --- | --- | --- | --- | --- |
| Treatment (between columns) | 102.7 | 2 | 51.35 | F (2, 24) = 8.349 |
| Residual (within columns) | 147.6 | 24 | 6.150 |  |
| Total | 250.3 | 26 |  |  |

| Tukey's multiple comparisons test | Mean diff. | 95.00% CI of diff. | Below threshold? | Adjusted P Value |
| --- | --- | --- | --- | --- |
| CONTROL vs. DREADD Gi | -4.750 | -7.759 to -1.741 | Yes | 0.0017 |
| CONTROL vs. FLOXED | -1.450 | -4.388 to 1.488 | No | 0.4461 |
| DREADD Gi vs. FLOXED | 3.300 | 0.4545 to 6.146 | Yes | 0.0209 |

**Dorsomedial hypothalamus (dmHyp)**
|  |  |  |  |  |
| --- | --- | --- | --- | --- |
| ANOVA table | SS | DF | MS | F (DFn, DFd) |
| Treatment (between columns) | 152.0 | 2 | 76.01 | F (2, 22) = 36.49 |
| Residual (within columns) | 45.82 | 22 | 2.083 |  |
| Total | 197.8 | 24 |  |  |
| Tukey's multiple comparisons test | Mean diff. | 95.00% CI of diff. | Below threshold? | Adjusted P Value |
| CONTROL vs. DREADD Gi | 6.278 | 4.367 to 8.189 | Yes | <0.0001 |
| CONTROL vs. FLOXED | 2.467 | 0.5945 to 4.339 | Yes | 0.0086 |
| DREADD Gi vs. FLOXED | -3.811 | -5.477 to -2.145 | Yes | <0.0001 |

**Dorsal periacueductal gray (dPAG)**
|  |  |  |  |  |
| --- | --- | --- | --- | --- |
| ANOVA table | SS | DF | MS | F (DFn, DFd) |
| Treatment (between columns) | 133.6 | 2 | 66.81 | F (2, 22) = 28.63 |
| Residual (within columns) | 51.33 | 22 | 2.333 |  |
| Total | 185.0 | 24 |  |  |
| Tukey's multiple comparisons test | Mean diff. | 95.00% CI of diff. | Below threshold? | Summary | Adjusted P Value |
| CONTROL vs. DREADD Gi | -5.667 | -7.689 to -3.644 | Yes | **** | <0.0001 |
| CONTROL vs. FLOXED | -1.667 | -3.648 to 0.3149 | No | ns | 0.1102 |
| DREADD Gi vs. FLOXED | 4.000 | 2.237 to 5.763 | Yes | **** | <0.0001 |

**Ventral periacueductal gray (vPAG)**
|  |  |  |  |  |
| --- | --- | --- | --- | --- |
| ANOVA table | SS | DF | MS | F (DFn, DFd) |
| Treatment (between columns) | 130.3 | 2 | 65.13 | F (2, 29) = 42.25 |
| Residual (within columns) | 44.71 | 29 | 1.542 |  |
| Total | 175.0 | 31 |  |  |
| Tukey's multiple comparisons test | Mean diff. | 95.00% CI of diff. | Below threshold? | Summary | Adjusted P Value |
| CONTROL vs. DREADD Gi | 3.208 | 1.809 to 4.608 | Yes | **** | <0.0001 |
| CONTROL vs. FLOXED | 5.208 | 3.809 to 6.608 | Yes | **** | <0.0001 |
| DREADD Gi vs. FLOXED | 2.000 | 0.7481 to 3.252 | Yes | ** | 0.0013 |

